# Hierarchical systems in the default mode network when reasoning about self and other mental states

**DOI:** 10.1101/2025.06.23.661113

**Authors:** Isaac R. Christian, Samuel A. Nastase

## Abstract

Humans spend time contemplating the minds of others. But this ability is not limited to external agents – we also turn the lens for reading minds inward, reflecting on our own thoughts, emotions, and sense of self. Some processes involved in reasoning about minds may rely on shared mechanisms, while others may be specific to the agent under consideration. We developed a paradigm where participants performed either a mental state inference task or a control task targeting either another person presented onscreen or their own mind. Using fMRI and multi-voxel pattern analysis, we replicate a well-established self-other axis along the medial wall of prefrontal cortex: ventral regions selectively decoded mental state inference patterns for self, but not other, whereas more dorsal regions decoded mental state inference for both self and other, compared to control conditions. Posterior cingulate cortex, on the other hand, differentiated the target of mental state inference. Using a cross-classification analysis, we also found patterns in the dorsomedial prefrontal cortex, ventromedial prefrontal cortex, and right temporoparietal junction were sensitive to mental state reasoning in general, regardless of the target agent. These findings highlight one process reflecting reasoning specific to the agent and another reflecting the reasoning process itself.

## Introduction

Social reasoning necessitates a theory of mind – flexible knowledge about the rich internal mental lives of agentic individuals (Ho, Saxe, & Cushman, 2022). Mental lives consist of, but are not limited to, beliefs, emotions, goals, and perspectives. We can reason about others’ beliefs, contemplating whether their current understanding of the world is true (Baron-Cohen, Leslie, & Frith, 1985; Wimmer & Perner, 1983). We can reason about their emotional states, deciding if they are upset (Lucas et al., 2014; Schmitgen, Walter, Drost, Rückl, & Schnell, 2016; Wu, Baker, Tenenbaum, & Schulz, 2018). We can reason about goals, determining if they intend to cooperate or compete (Camerer, Ho, & Chong, 2004; Lee & Seo, 2016). We can effortlessly take the perspective of friends and celebrities, predicting their view in a range of contexts (Batson, Early, & Salvarani, 1997).

At the core of these abilities lies mental state inference (Mitchell, 2009). Computationally, mental state inference is defined as updating prior beliefs and goals about an agent when new evidence becomes available (Baker, Saxe, & Tenenbaum, 2011; Charpentier & O’Doherty, 2018; Jara-Ettinger, 2019). More generally, inference can be understood as the strategic assembly of available cues, both observable and unobservable, to predict a hidden variable – in this case a mental state (Frith & Frith, 1999; Tamir & Thornton, 2018). The type of inference depends on the nature of the task and social context (Ho et al., 2022; Jara-Ettinger & Dunham, 2024; Ullman & Bass, 2024). If required to estimate an agents’ current mental state, we may rely on a combination of external cues such as eye gaze, posture, or their actions. If tasked with predicting how a friend will act in a new situation, we might leverage a mental model of that person developed from previous experience, or use our own personal experience as a reference.

Just as we can reason about other individuals, so too can we reason about our own minds. We introspect, considering our own dynamic perceptions, emotions, intentions, and beliefs, sometimes asking ‘How am I feeling?’ or ‘Am I extroverted?’. This ability to ascribe a second order representation of internal states and processes is called metacognition (Fleming, Dolan, & Frith, 2012). The process of representing one’s own mind and the minds of others may be quite similar (Graziano, 2013; Heyes, Bang, Shea, Frith, & Fleming, 2020; Vaccaro & Fleming, 2018). In fact, it has been proposed that metacognition is merely turning the lens for reading others’ minds on oneself (Carruthers, 2009; Frith, 2012). This theory suggests that similar mechanisms, including processes for belief inference and action representation are deployed irrespective of agent type (Gallese & Goldman, 1998). Other mechanisms, in contrast, may be unique to the agent under consideration. When reasoning about our own mental states, we rely heavily on internal cues, such as interoceptive signals (e.g. heart rate, arousal) and memory to infer a mental state (e.g. I am feeling excited) rather than external cues (Picard & Friston, 2014; Seth, 2013). Thus, while some social reasoning processes may generalize across agent types, others may be specific to the self or other agents.

Social reasoning is associated with a network of regions overlapping the default mode network (DMN; Andrews-Hanna, Reidler, Sepulcre, Poulin, & Buckner, 2010; Mars et al., 2012). This network appears to be specialized for social reasoning, with a separate network associated with reasoning more broadly (Van Overwalle, 2011). Primary nodes of the DMN include the dorsal medial prefrontal cortex (DMPFC), the ventral medial prefrontal cortex (VMPFC), the posterior cingulate cortex (PCC), and left and right temporoparietal junction (LTPJ and RTPJ). These components may have distinct roles that reflect agent-specific or agent-general reasoning. The DMPFC, for instance, is sometimes implicated in computing an inference that generalizes across agents (Nicolle et al., 2012; Wittmann et al., 2016; Yoshida, Seymour, Friston, & Dolan, 2010). Other studies, however, find the DMPFC is functionally tuned to the processing of others, and is embedded in an agent-specific representational hierarchy along the medial wall of prefrontal cortex (Amodio & Frith, 2006; Bzdok et al., 2013; Jamali et al., 2021; Mitchell, Macrae, & Banaji, 2006; Sul et al., 2015; Wagner, Haxby, & Heatherton, 2012). In this account, the DMPFC is primarily engaged when reasoning about others, activating during inferences about other agents’ beliefs (Saxe & Kanwisher, 2013), strategies (Coricelli & Nagel, 2009), emotional states (Zaki, Weber, Bolger, & Ochsner, 2009), or their level of expertise (Boorman, O’Doherty, Adolphs, & Rangel, 2013). In contrast, the VMPFC exhibits sensitivity to self-referential information (Northoff et al., 2006; Rogers, Kuiper, & Kirker, 1977), responding when considering one’s own personality (Macrae, Moran, Heatherton, Banfield, & Kelley, 2004), affective state (Gusnard, Akbudak, Shulman, & Raichle, 2001; Y. Suzuki & Tanaka, 2021), and when adopting a first person perspective (Vogeley et al., 2004). Self-other processing appears to be modulated by the perceived similarity between oneself and another agent: thinking about similar others elicits VMPFC activity, while reasoning about dissimilar others engages more dorsal regions (Aron, Aron, Tudor, & Nelson, 1991; Krienen, Tu, & Buckner, 2010; Mitchell et al., 2006).

The present study investigated both general and agent-specific representations involved in reasoning about a mental state. To this end, we designed a novel functional magnetic resonance imaging (fMRI) paradigm that required participants to infer either their own mental state or the mental state of another agent performing the same task as depicted in a video stimulus. Our study consisted of four conditions, two tasks that involved either reasoning about the mental states of other (Other Reason) or self (Self Reason) agents and two control conditions that did not involve explicit mental state inference (Self Count, Other Count). In the Other Reason task, participants were asked to infer the content of another’s mind – whether or not an agent had ‘mind wandered’ from the task – using cues available in the video stimulus. In the Self Reason task, participants made an inference about the content of their own mind – whether or not their attention had wandered from the task – using cues available in their mind. In control conditions, participants were asked to perform a counting task that depended on another agent’s breathing sequence or their own breathing sequence. These control conditions still required the use of cues – for example, the physiological sensation of respiration (Self), or observation of the rise and fall of the other agent’s chest during respiration (Other) – but without the need to use these cues for mental state inference.

We hypothesized that the self/other distinction and general mechanisms for inference would be hierarchically organized in the DMN. We used multi-voxel pattern analysis to explore these hypotheses, predicting greater sensitivity in the VMPFC to self-reasoning compared to control conditions, and greater sensitivity in the DMPFC to other-reasoning compared to control conditions. We then conducted analyses to identify processes that were generally associated with social reasoning or were specific to agent type.

## Methods

Twenty-eight healthy human volunteers (aged 18-50, normal or corrected to normal vision, 27 right handed, 16 female) were recruited from the community and from a subject pool sponsored by Princeton University. All subjects provided consent and received either $40 or course credit for participation. All procedures were approved by the Princeton Institutional Review Board.

The task design, shown in Figure 1, involved four conditions: Self Reason, Self Count, Other Reason, and Other Count. In the Self Reason condition, subjects were instructed to pay attention to the rhythmic sensation of their breathing for five minutes. If at any point during this period they noticed that their mind had wandered from their breath, described as ‘mind wandering’, they were to indicate the moment by pressing the button, and then reorient attention back to their breath and continue the task. The period right before the button press, during which participants inspected their current mental state, is of greatest interest in the present study. We hypothesized that this time period represents the most intensive mentalization about one’s own mind, and that participants would infer whether their current attention state matched a definition of ‘on task’ at that moment. We extracted activity from this period (–3 to 0 s) before the button press. The moment of realization presumably took less than three seconds, but we used a broad 3 s window to capture variation in the mentalization process according to previous work (Hasenkamp, Wilson-Mendenhall, Duncan, & Barsalou, 2012).

**Figure 1:**
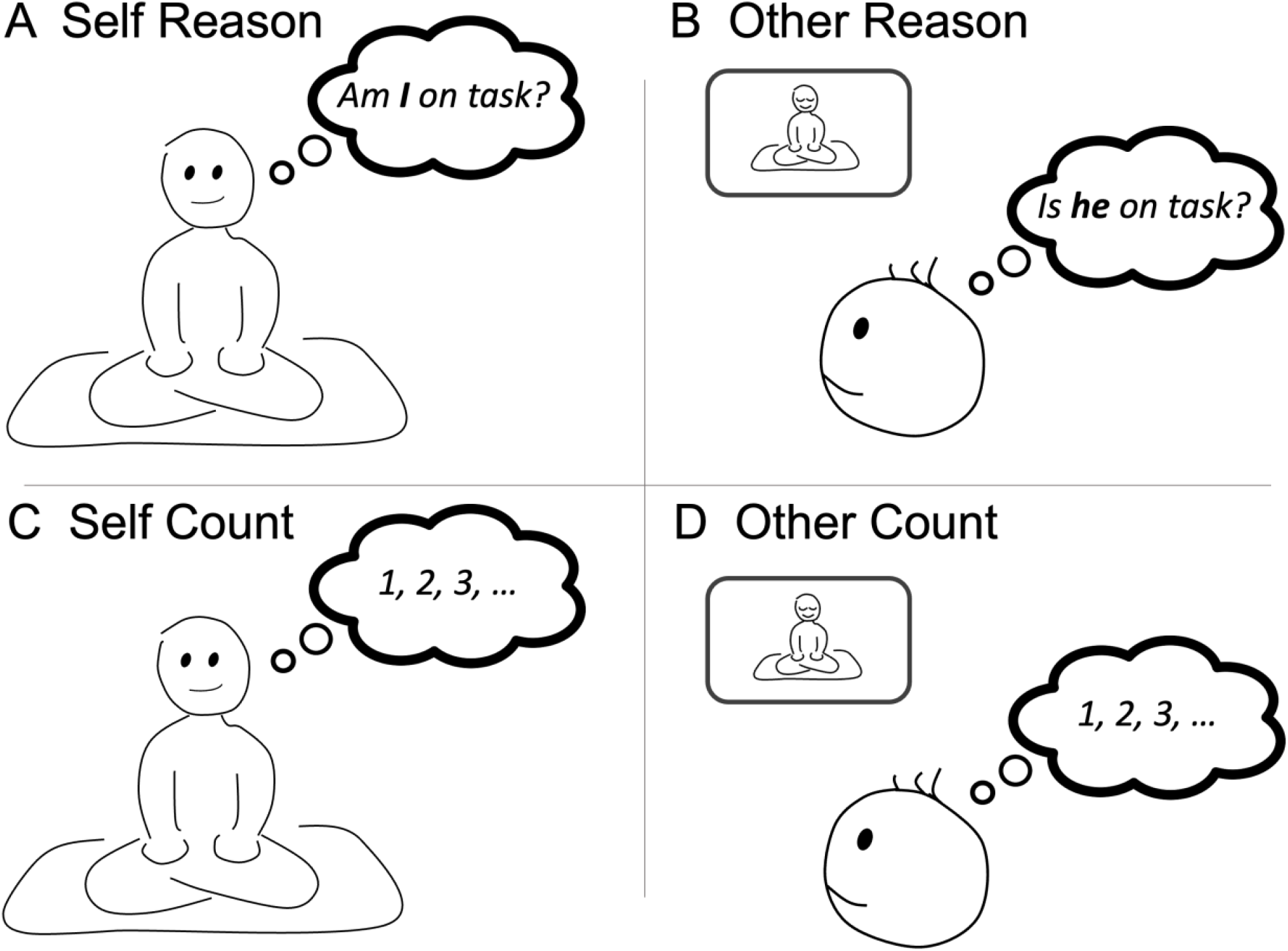
Self/other reasoning and control tasks. (A) In the Self Reason task, subjects monitored the internal content of their mind and decided when their mental state was no longer on task. (B) In the Other Reason task, subjects monitored the mental state of an agent in a video, deciding when they believed the agent was no longer on task. (C) In the Self Count task, subjects counted their own breaths and pressed a button at every fifth breath. (D) In the Other Count task, subjects counted the breaths of an agent in a video and pressed a button at every fifth breath.

In the Other Reason condition, participants assessed the content of another mind by watching a movie with an agent performing the Self Reason task. The film showed the agent from the waist up, revealing only the upper body and face with eyes closed. Subjects were asked to carefully attend to the agent in the movie, to try to assess on the basis of any cues (body posture, facial expression, and so on) whether the agent had suffered a moment of mind wandering, and to press the button at those moments of perceived mind wandering. We hypothesized that a mechanism involved in constructing models of someone else’s mental state should show activity in this period.

In addition to the Self Reason and Other Reason conditions, we included two control conditions. These conditions were intended to incorporate many of the same cognitive processes as the Reasoning tasks, such as attention on breath, focused control of attention, and a button press response, but they were designed to exclude the crucial moment when subjects determined whether their own mental state or the mental state of another agent was on task. For the Self Count task (Figure 1C), subjects once again focused on their own breathing. They were instructed to count breaths, and on reaching a count of five, to press the button and begin the count again at 1. For the Other Count condition (Figure 1D), subjects were instructed to count the breaths of the agent in the video, and, on reaching a count of five, to press the button and to begin the count again at 1.

To make the stimulus videos for the Other Reason and Other Count conditions, two volunteer agents were recorded (agent A and B), for two separate five-minute films. The agents sat cross legged on a cushion and rested the right index finger on the spacebar of a computer used to record button presses. The agents were told to focus their attention on their own breathing and to press the button at the moment they realized their mind had wandered from the task. The camera was positioned to exclude view of the hands and computer (and thus of the button presses) and to capture an image of the actor above the waist.

Each subject performed 10 runs of 5 minutes each. First, the subject performed four runs corresponding to the four behavioral conditions in a randomized order. The same video, showing agent A, was presented for both the Other Reason and Other Count conditions. The subject then performed a second iteration of the four runs, corresponding to the four behavioral conditions in a different randomized order. For this second iteration, the video of agent B was presented for the Other Reason and Other Count conditions. In addition to the eight runs in which the behavioral task was performed, subjects performed two, five-minute runs of rest, one at the start of testing and one at the end. Between runs, participants were offered a 30-s rest that they could skip by pressing a “continue” button. Total scan time was approximately one hour.

### fMRI acquisition and processing

Functional imaging data were collected using a 3T MAGNETOM Skyra scanner equipped with a 64-channel head/neck coil. Gradient-echo T2*-weighted echo-planar images (EPI) with blood oxygen level dependent (BOLD) contrast were used as an index of brain activity: TR = 1.5 s, voxel size = 2.5 × 2.5 × 3 mm. Data were preprocessed using fMRIPrep version 20.2.0. T1-weighted volumes were skull-stripped using the OASIS template in antsBrainExtraction.sh v2.1.0. Spatial normalization through nonlinear registration to the ICBM 152 Nonlinear Asymmetrical template version 2009c was completed using the antsRegistration tool of ANTs v2.1.0 (Avants, Epstein, Grossman, & Gee, 2008). Brain tissue segmentation of cerebrospinal fluid (CSF), white matter (WM) and gray matter (GM) using FAST was performed on extracted T1w images.

Functional data were slice time corrected using 3dTshift from AFNI v16.2.07 and motion corrected using MCFLIRT (FSL v5.0.9). FLIRT (FSL) performed boundary-based registration with six degrees of freedom to co-register the corresponding T1w images to functional data. Motion correcting transformations, BOLD-to-T1w transformation, and T1w-to-template warp were concatenated and applied in a single step using antsApplyTransforms (ANTs v2.1.0) with Lanczos interpolation. All functional images were low-passed (0.25) using Nilearn’s signal cleaning function. Physiological noise regressors included the first 5 principle components for CSF and WM for each functional run, totaling 10 aCompCor components. Head motion parameters included 3 translation and 3 rotation time series as well as censor time series for volumes with a framewise displacement (FD) exceeding 0.3 mm. For each volume with FD exceeding 0.3 mm a vector of zeros was constructed with a value of one assigned to the time point corresponding to the offending volume. If more than 30% of volumes in a run had an average FD of 0.3 mm or larger, those runs were omitted from the analysis (9.6% of runs met this stringent criterion). For each run, censor time series as well as 10 aCompcor components, 6 head motion parameters and 3 cosine drift parameters were inserted as regressors of no interest into the subsequent General Linear Models (GLMs).

A separate GLM was constructed for each button press event. To set up the GLM, we defined a period of interest as 3 to 0 s prior to the button press for each of four behavioral conditions (Self Reason, Self Count, Other Reason, Other Count). The predicted BOLD activity for each 3 s period prior to the button press was treated as a regressor of interest in a single design matrix along with the nuisance regressors. First-level regression was performed using Nilearn’s FirstLevelModel function. Regressors of interest were convolved with a canonical hemodynamic response function (Glover, 1999). Regression coefficients (beta weights) for each button press were extracted from the resulting images and used in multivariate analysis. The BOLD data were spatially smoothed using a 2 mm full width half maximum Gaussian kernel prior to regression to facilitate pattern classification across subjects.

## Multi-voxel pattern analysis

We selected five target regions of interest (ROIs) from five primary nodes of the DMN. These regions included the DMPFC, VMPFC, RTPJ, LTPJ, and the PCC. These regions were selected from a search of ‘mentalizing’ in Neurosynth (Yarkoni, Poldrack, Nichols, Van Essen, & Wager, 2011) which yielded an averaged activation map across 151 studies that used the term. From this map, we created a 10 mm sphere using the maximum t-statistic as the center for each target region, and used these ROI masks to extract beta weights from each subject’s whole brain image from the univariate analysis.

A linear support vector classifier (SVC) was trained on one subset of subjects and tested on a left-out subset of subjects using scikit-learn (https://scikit-learn.org/). Because some subjects pressed the button relatively few times per condition, we did not use a leave-one-subject-out procedure, which would have left too few samples for the test set in some cases. Instead, we used 5-fold leave-one-group-out cross-validation across subjects. Subjects were first split into five groups (5 subjects in four of the groups and 8 subjects in one group). We trained the SVC on four groups and tested on one group, optimizing the parameters for the model using nested cross-validation: within each training set, we performed a grid search across the SVC regularization parameters C ranging from 10^−5^ to 10 using 5-fold leave-one-group-out cross-validation, and retrained the best performing model to predict the outer test set. Ultimately, this procedure resulted in five classification accuracies, which were averaged to produce the final accuracy score. To evaluate the statistical significance of that accuracy score, we created a null distribution by running 10,000 iterations of the SVC with randomly shuffled condition labels. The true decoding accuracy for each analysis was compared to the permutation-based null distribution to obtain a p-value. We additionally defined a 95% confidence interval around the accuracy score by bootstrapping the five test statistics obtained from each fold during cross validation. These five values were resampled with replacement and their means calculated a total of 10,000 times. The upper (97.5%) and lower (2.5%) percentiles of the bootstrap means were then computed to establish confidence intervals.

This procedure was used for five analyses. Two of these analyses compared the voxel patterns of target reasoning conditions to the voxel patterns of control conditions (Self Reason compared to Self Count; Other Reason compared to Other Count). In the third test, we directly compared the voxel patterns of reasoning conditions (Self Reason compared to Other Reason). In the fourth test, we compared the voxel patterns of both control conditions (Self Count compared to Other Count).

In the final test, we trained a classifier on the distinction between the Self Reason and Self Count conditions, and then tested whether, once trained, the classifier could decode the difference between Other Reason and Other Count. This cross-classification procedure (Kaplan et al., 2015) allowed us to test whether an ROI differentiates the Reason versus Count conditions in a manner that generalizes across Self and Other conditions.

## Results

Across subjects, the mean number of button presses during a five minute run, for each of the four tasks, was: Self Reason, mean = 10.86, SEM = 0.81 ; Self Count, M = 12.05, SEM = 0.53; Other Reason, M = 11.16, SEM = 1.00; and Other Count, M = 11.75, SEM = 0.39. Button press counts did not significantly differ between Reason and Count conditions or between Self and Other conditions (2 × 2 within subjects ANOVA: for Reason versus Count conditions, F = 0.78, p = 0.38; for Self versus Other conditions, F = 0.08, p = 0.78; for interaction, F = 0.14, p = 0.71).

We performed five decoding analyses with five ROIs in the DMN: the DMPFC, VMPFC, PCC, the right and left TPJ, (Figure 2F).

First, we tested whether the pattern of activity within an ROI differentiated the Self Reason task from the Self Count task (Figure 2A) and whether the pattern of activity for the Other Reason task differed from the Other Count task (Figure 2B). In the Self Reason versus Self Count comparison, patterns within the DMPFC and VMPFC were distinct, at a peak accuracy of 55.3% (significantly above chance based on a permutation test, *p* < 0.01) for the DMPFC and 54.2% (*p* < 0.05) for the VMPFC. When classifying Other Reason versus Other Count, only the DMPFC showed significant decoding at 56.3% (*p* < 0.001).

**Figure 2:**
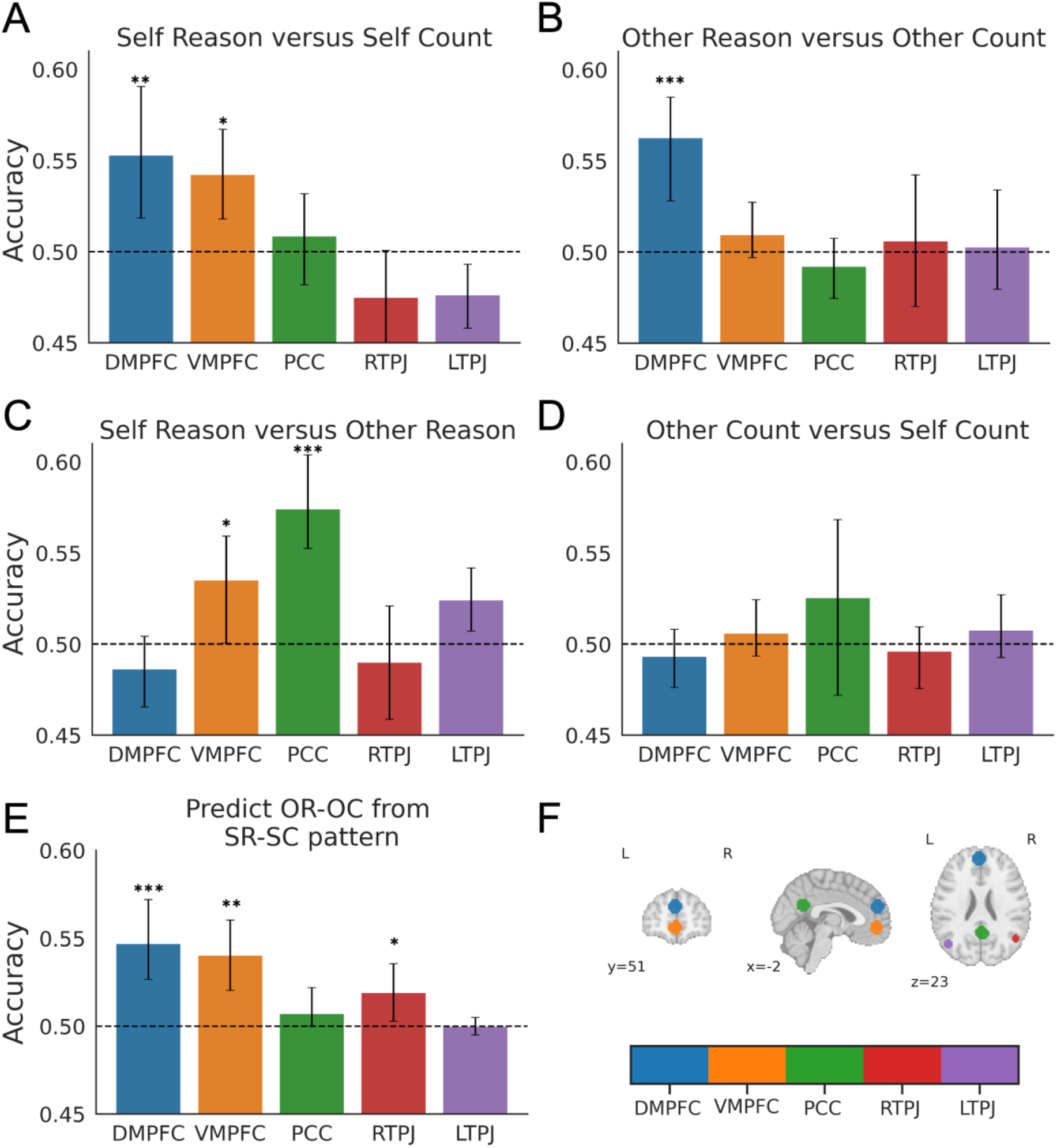
Decoding neural representations for self/other reasoning. (A) Classification results for Self Reason versus Self Count tasks across five ROIs associated with mentalizing. Activity patterns correspond to the 3 s period during which participants reasoned if they were on task or if their breath count had reached five. Dotted line shows chance accuracy of 50%. Statistical significance was assessed using a permutation test (*** *p* < 0.001, ** *p* < 0.01, * *p* < 0.05). Error bars depict 95% bootstrap confidence intervals around the mean classification accuracy. (B) Classification results for Other Reason versus Other Count tasks. (C) Classification results for Self Reason versus Other Reason tasks. (D) Classification results for Other Count versus Self Count tasks. (E) Cross-classification results when training the classifier to distinguish activity patterns for Self Reason versus Self Count tasks, and then testing the generalization of the trained classifier to the Other Reason versus Other Count tasks. (F) Five ROIs in the DMN associated with mentalizing.

These results show that there is agent specificity when reasoning tasks are compared to control conditions. Other regions in the DMN, however, may track the agent targeted during reasoning. To test this hypothesis, we directly compared Self Reason and Other Reason conditions. We observed significant classification accuracy in the PCC (accuracy = 57.4%, p < 0.001), a region of the DMN not implicated in the prior analyses (Figure 2C).

In a control analysis, we trained a classifier to differentiate the Count conditions, and found that none of the DMN regions were able to distinguish self versus other in the Count tasks. This suggests that the preceding decoding results are driven by mental state reasoning: patterns in the DMN distinguished self versus other processing when reasoning about mental states, but not when counting breaths.

Given that activity patterns in DMPFC appeared to differentiate Reason and Count tasks for both Self and Other conditions, we used a cross-classification analysis to more rigorously test if the patterns in DMPFC reflect social reasoning in a way that generalizes across the agent under consideration (self or other). A significant accuracy would suggest that DMPFC differentiates social reasoning from counting in a similar way, whether reasoning about one’s own mental states or those of another agent. As shown in Figure 2E, the decoding is significant for the DMPFC, with a peak accuracy of 54.7% (*p* < 0.01). For completeness, we also performed cross-decoding for the other four ROIs. As shown in Figure 2E, cross-classification was also significant in the VMPFC (accuracy = 54.0%, p < 0.001) and the RTPJ (accuracy = 51.92%, p < 0.05).

## Discussion

We assessed neural systems underlying reasoning about one’s own and another’s mental state. We found that patterns in the DMPFC, VMPFC and PCC were sensitive to the type of agent involved (Fig. 2A, 2B, 2C), and that patterns in the DMPFC, VMPFC, and RTPJ predicted mental state inference irrespective of the agent in question (Fig. 2E). These findings reveal neural processes for social reasoning that are both general and specific to the agent under consideration.

We replicated a processing hierarchy sensitive to the kind of agent – self or other – engaged in social reasoning (Amodio & Frith, 2006; Bzdok et al., 2013; Denny, Kober, Wager, & Ochsner, 2012). Note that our across-subject decoding analysis focused on distinguishing activity patterns across individuals. A within-subject decoding analysis may better accommodate individual differences in functional topography. However, our participants only pressed the button on average ∼11 times per run. Given relatively few samples within subjects, we opted for across-subject pattern analysis (e.g., Poldrack et al., 2009; Chen et al., 2017; Zadbood et al., 2018). This finding supports prior work suggesting a self-other axis, including paradigms examining value-based decision making (Ereira, Dolan, & Kurth-Nelson, 2018; Sul et al., 2015), social knowledge (Mitchell, 2009; Wagner et al., 2012), and competitive gaming (Gallagher, Jack, Roepstorff, & Frith, 2002). Many of these studies rely on comparisons between target conditions (self, other) and control conditions, similar to our comparison between reasoning and counting conditions. However, results regarding a self-other axis are less consistent when self and other conditions are directly compared. One study reports no significant differences in univariate responses between self and other conditions (Vanderwal, Hunyadi, Grupe, Connors, & Schultz, 2008), while others find greater DMPFC activation for self compared to other (Benoit, Gilbert, Volle, & Burgess, 2010; D’Argembeau et al., 2007; Krienen et al., 2010).

When we directly compared self and other reasoning conditions, we found that patterns in the VMPFC and the PCC identified which agent was being reasoned about. In contrast to the VMPFC, which showed a sensitivity in other comparisons, the PCC only distinguished patterns when self and other reasoning were directly compared. This agent-specific processing may stem from the PCC’s role in integrating past experiences, semantic knowledge, and current perceptual information (Cavanna & Trimble, 2006; Sugiura, Shah, Zilles, & Fink, 2005; Yeshurun, Nguyen, & Hasson, 2021). Furthermore, although the PCC typically exhibits a heightened sensitivity for self-related processing (Davey, Pujol, & Harrison, 2016; Moran, Lee, & Gabrieli, 2011), the PCC (along with the other DMN nodes tested) did not distinguish between agents when the task did not involve reasoning, but agents were directly compared. This finding suggests the PCC is not more sensitive to one agent over the other or picking up on low-level feature differences between agents (e.g. internal or external cues, visual information). Instead, the PCC appears to differentiate agents specifically during higher-order cognitive processes like social reasoning, potentially updating mental models that are agent specific.

Patterns in the DMPFC were sensitive to reasoning about oneself and another agent. The DMPFC is well known to play a role in other-oriented processing (Fletcher et al., 1995; Goel, Grafman, Sadato, & Hallett, 1995), but less so in self-referential processing (although see exceptions: Benoit et al., 2010; Frith & Frith, 1999; Wittmann et al., 2016). Many paradigms may miss the DMPFC’s involvement in self processing because they fail to fully engage a self-related inference. Comparing personality traits to self-concepts, as is standard in research on self-other processing, elicits semantic comparison (i.e. comparing one’s representation of self to a representation of a character trait) rather than inferential reasoning. Furthermore, classic Theory of Mind paradigms, such as false-belief tasks, although useful for targeting other-related reasoning, have no clear analog for reasoning about one’s own mental state. Our task, in contrast, may capture an inference about one’s current mental state, as we explicitly asked participants to use the internal evidence available to hypothesize an unknown variable (i.e. on task or off task).

We acknowledge that this definition of inference is not as precise as standard computational formulations (e.g. Collette, Pauli, Bossaerts, & O’Doherty, 2017). However, these approaches often do not measure inference that is directed towards one’s own mind per se, instead defining self-oriented processing in the context of reinforcement learning, as actions made for oneself (Nicolle et al., 2012) or one’s own value preferences (Campbell-Meiklejohn, Simonsen, Frith, & Daw, 2017; Garvert, Moutoussis, Kurth-Nelson, Behrens, & Dolan, 2015), for instance. We believe our task is a step forward in characterizing a similar inference procedure for both self and other agents, and our findings indicate that DMPFC’s role in reasoning about the mental states of others may extend to reasoning about our own mental states.

We found that representations in DMPFC and VMPFC encode information relevant for inferring mental states across agents (Fig. 2E), even though they exhibit a nuanced ability for individual agents, as discussed above. One explanation for this result is that both social- and meta-cognitive processes rely on shared capacity for abstraction (Vaccaro & Fleming, 2018). Our task requires participants to explicitly (as opposed to implicitly) evaluate and track task performance of their own and another agent’s mental state. The observed generalizability of VMPFC and DMPFC across agents may therefore reflect a broader mechanism involved in abstracting, rather than a function exclusive to social processing, like mental state inference.

Lastly, the RTPJ (in addition to the VMPFC) showed generalizability across agent type, despite no decoding accuracy when discerning activation patterns between Self Reason and Self Count or Other Reason and Other Count. This may seem confusing – how can representations generalize for social reasoning across agents, yet fail to show significance in the more basic reasoning and counting comparisons for either agent? One possible explanation is that the cross-classification analysis was better powered to detect differences in the signal compared to the standard classification analyses. Alternatively, certain features shared between conditions (e.g., other reasoning versus other count) may obscure features of the activity used to differentiate reasoning versus count in the cross-classification analysis. Furthermore, describing one’s own mental state and the mental state of another is a highly unconstrained process, with events occurring at different points in the task and the heuristics varying across participants and trial contexts. To solidify the RTPJ’s role in agent-general reasoning, future studies should utilize a larger sample size to determine whether the cross-classification results persist in the absence of within-condition classification.

In sum, we use a novel task to characterize the modular components involved in agent specific and agent general processing, finding that some regions of the DMN are sensitive to agent type and social reasoning more generally, while others distinguish reasoning targeting the self and others. We interpret our results to support a two-level hierarchical structure where the lower-order processes track mental states in an actor-specific way, while the higher-order processes are more abstract, generalizing mental state inference across actors (self or other).

## Notes

**Conflict of Interest:** The authors declare no conflict of interest.

**Funding / Acknowledgements:** Supported by Princeton Neuroscience Institute Innovation Fund 24400E2349FA010 and by AE Studios grant 24400B1459FA010.

### Competing Interest Statement

The authors have declared no competing interest.

